# Comparative thermal performance of *Orbicella franksi* at its latitudinal range limits

**DOI:** 10.1101/583294

**Authors:** Nyssa J. Silbiger, Gretchen Goodbody-Gringley, John F. Bruno, Hollie M. Putnam

## Abstract

Temperature drives biological responses that scale from the cellular to ecosystem levels and thermal sensitivity will shape organismal functions and population dynamics as the world warms. Reef building corals are sensitive to temperature due to their endosymbiotic relationship with single celled dinoflagellates, with mass mortality events increasing in frequency and magnitude. The purpose of this study was to quantify the thermal sensitivity of important physiological functions of a Caribbean reef-building coral, *Orbicella franksi* through the measurement of thermal performance curves (TPCs). We compared TPC metrics (thermal optimum, critical maximum, activation energy, deactivation energy, and rate at a standardized temperature) between two populations at the northern and southern extent of the geographic range of this species. We further compared essential coral organismal processes (gross photosynthesis, respiration, and calcification) within a site to determine which function is most sensitive to thermal stress using a hierarchical Bayesian modeling approach. We found evidence for differences in thermal performance, which could be due to thermal adaptation or acclimatization, with higher TPC metrics (thermal optimum and critical maximum) in warmer Panama, compared to cooler Bermuda. We also documented the hierarchy in thermal sensitivity of essential organismal functions within a population, with respiration less sensitive than photosynthesis, which was less sensitive than calcification. Understanding thermal performance of corals is essential for projecting coral reef futures, given that key biological functions necessary to sustain coral reef ecosystems are thermally-mediated.

**Summary statement:** We apply a thermal performance curve approach to a variety of fitness related parameters in a reef building coral across its geographic range and various functions to improve our understanding of the inherent variability in thermal tolerance.

## Introduction

Evidence that anthropogenic climate change is impacting the natural world continues to accumulate (Parmesan and Yohe, 2003; Poloczanska et al., 2013). Populations in all types of habitats — aquatic and terrestrial, from the poles to the equator — are responding to warming (Oliver et al., 2018; Walther et al., 2002) and countless other important environmental changes (U S Global Change Research Program, 2019). Essential to improving predictions of organismal to ecosystem responses to environmental change, is the quantification of not only individual/genotype and population response to warming, but also functional sensitivities.

The sensitivity of ecotherms to temperature can be characterized empirically as a Thermal Performance Curve (TPC), which quantifies the shape of the relationship (typically following a Sharpe-Schoolfield model; (Schoolfield et al., 1981; Sharpe and DeMichele, 1977)) between biological rates of “performance” (e.g., respiration or growth) and environmental temperature (Figure 1; Table 1). TPC parameters such as critical minimum (CT_min_), critical maximum (CT_max_), and thermal optimum (T_opt_) describe the limits and optima of a chosen process with changing temperature (Angilletta, 2009; Huey and Kingsolver, 1989; Huey and Stevenson, 1979). The positive and negative slopes on either side of T_opt_ are the rates of activation (E) and deactivation (E_h_) energy, respectively, and indicate the sensitivity of the process. Species with high E and E_h_ (i.e. steeper slopes on either end of the curve) will be most sensitive because they will quickly move from optimal to suboptimal conditions with only small changes in temperature. The rate at a standardized temperature, b(T_c_), is often used to compare rates between organisms or functions at a reference temperature (Padfield et al., 2017). Each of these TPC parameters can be compared among populations and species, locations with different thermal histories (e.g., across latitudes), and over time to predict future community composition and the functional consequences of environmental change.

**Table 1:** Equations and parameters used in constructing thermal performance curves and calculating derived quantities.

| Model/Derived Quantities | Equation |
| --- | --- |
| Modified Sharpe-Schoolfield for high temperature inactivation | $\log(\text{rate}) = b(T_c) + E\left(\frac{1}{T_c} - \frac{1}{k \cdot T_i}\right) - \log\left(1 + e^{Eh\left(\frac{1}{K \cdot Th} - \frac{1}{K \cdot T_i}\right)}\right)$ |
| Thermal optimum | $T_{opt} = \left(\frac{Eh \cdot Th}{Eh + (K \cdot Th_k \cdot \log(\frac{Eh}{E} - 1))}\right)$ |
| Parameters | Description |
| $b(T_c)$ | Log rate at a constant temperature ($\mu\text{mol cm}^{-2} \text{ hr}^{-1}$) |
| $E$ | Activation energy (electron volts, eV) |
| $Eh$ | Temperature-induced inactivation of enzyme kinetics past $Th$ for each population (electron volts, eV) |
| $K$ | Boltzmann constant ($8.62 \times 10^{-5} \text{ eV K}^{-1}$) |
| $T_c$ | $T_c$ is the reference temperature at which no low or high temperature inactivation is experienced (defined here as 300.15 K or 27°C) |
| $T_h$ | Temperature (K) at which half the enzymes are inactivated |
| $T_i$ | Temperature $i$ in Kelvin (K) |

**Figure 1.**
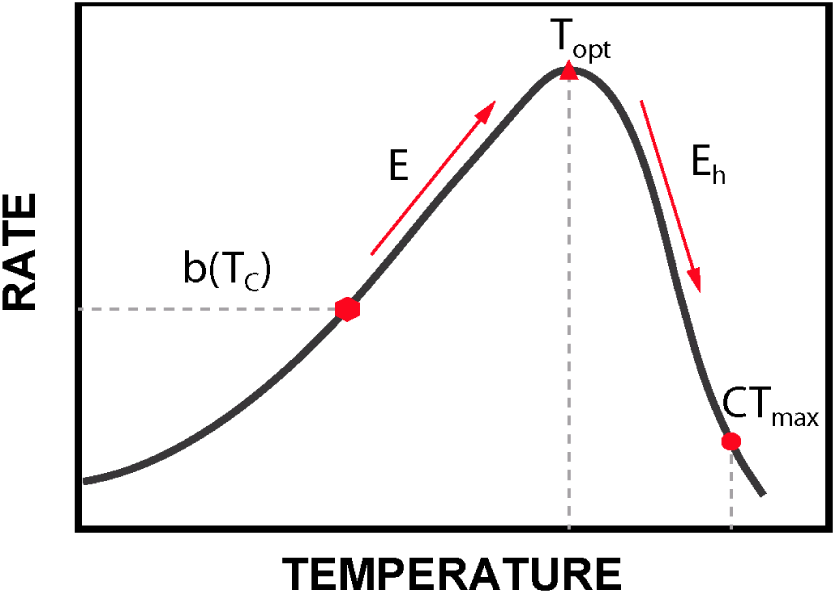
Thermal performance curve characteristics, with comparative temperature metrics identified by the red symbols.

In the context of climate change, applying a TPC approach to quantify thermal adaptation is essentially a space-for-time substitution (Blois et al., 2013; Faber et al., 2018; Pickett, 1989) — a widely used approach when the practical temporal extent of a study cannot match the process of interest. There is extensive evidence of “adaptation” to spatial temperature gradients in a wide range of organisms, from local (e.g., tens of meters) to regional (100s of km) scales (Berkelmans, 2002; Berkelmans and Willis, 1999; Castillo et al., 2012; Oliver and Palumbi, 2011). Such phenotypic gradients presumably reflect an underlying selection gradient and can provide clues about numerous aspects of thermal adaptation, such as fitness costs and other trade-offs and the plausible range of survivable temperatures.

Coral reefs are being severely affected by global warming. Like many other foundation species, the corals that build tropical reefs are being lost in response to anthropogenic warming (Hughes et al., 2017; Hughes et al., 2018b). Coral cover has declined dramatically on reefs worldwide over the last 30 years (from ~60% to <20% on some reefs), in large part due to ocean warming (Bruno and Selig, 2007; De’ath et al., 2012; Gardner et al., 2003; Hughes et al., 2018a) causing mass bleaching, or the loss of the corals’ symbiotic dinoflagellates and their nutritional benefits to the host (Oakley and Davy, 2018). While there are numerous local causes of coral loss (e.g., pollution, destructive fishing practices, tourism, etc.), the single most detrimental stressor to date is thermal stress from anomalous heating events (i.e., heatwaves) and its associated complications (i.e., bleaching, disease, reduced calcification etc.; (DeCarlo et al., 2017; Harvell et al., 2002; Hughes et al., 2017)). Reef building corals and their dinoflagellate symbionts live close to their physiological thermal maximum and, as a result, warming of 1°C or more above local mean monthly maxima can reduce fitness and cause tissue loss or whole-colony mortality (Hoegh-Guldberg 1999, Baker et al. 2008), with significant negative implications for reef structure and function (Couch et al., 2017; Hughes et al., 2018b; Stuart-Smith et al., 2018). In the case of corals, there are numerous mechanisms that could enable local thermal adaptation, including genetic adaptation and physiological acclimatization of the host, changes in Symbiodinaceae composition, and the bacterial microbiome (Putnam et al., 2017).

Corals have several key physiological traits that scale up to influence ecosystem function. These traits include production, respiration, and calcification, which generate much of the carbon on a coral dominated reef, power coral metabolic processes, and build the 3-dimensional structure of the reef, respectively. Importantly, these traits are likely to have different responses to temperature. For example, coral holobiont photosynthesis is dependent on single celled dinoflagellates (in the family Symbiodiniaceae; (LaJeunesse et al., 2018)). These endosymbionts act as light and temperature sensors for the coral host and typically initiate the cascade of dysbiosis and bleaching through the generation of reactive oxygen species production, due to excess excitation energy in the photosystems under increasing temperatures (Oakley and Davy, 2018). Additionally, different enzymatic machinery is utilized for these different traits, with likely differing thermal optima. Therefore, in comparison, holobiont respiration is likely to be less sensitive to temperature than photosynthesis. Further, because light-enhanced calcification is hypothesized to be dependent on energy supplied by Symbiodiniaceae (Allemand et al., 2011), corals may substantially reduce calcification when production is low (e.g., during bleaching events; (Barkley et al., 2018)). Lastly, the ratios between these different processes can change with temperature and have implications for survival and the long-term persistence of certain coral functions (Coles and Jokiel, 1977).

The purpose of this study was to quantify the thermal sensitivity of important physiological functions of a reef-building coral with the goals of 1) comparing responses among colonies (putative clones) of the Caribbean coral *Orbicella franksi*, between two populations at the northern (cool, Bermuda) and near the southern (warm, Panama) extent of the geographic range of this species and 2) comparing essential coral organismal processes (gross photosynthesis, respiration, and calcification) within a site (Bermuda) to determine which function is most sensitive to thermal stress. We hypothesized that corals in Panama would have higher thermal optima than Bermuda based on thermal history and that respiration rates would be more thermally tolerant than photosynthesis and calcification based on symbiont sensitivity to light and temperature and evidence for their contributions to dysbiosis (Lesser, 2011; Oakley and Davy, 2018; Venn et al., 2008). To test these hypotheses, we quantified TPCs using a set of hierarchical Bayesian models.

## Materials and Methods

### Study Sites (Panama and Bermuda)

The Bocas del Toro Archipelago is located on the Caribbean coast of Panama at 9°N, 82°W on the border of Costa Rica (Figure 2). It is composed of a complex network of islands and mainland peninsulas fringed by mangroves with well-developed seagrass beds and coral reefs (Collin, 2005). The region hosts a high diversity of scleractinian corals, with 61 species documented, and mean coral coverage of 26.9% (Guzman et al., 2005). Long-term temperature records from shallow fringing reef systems within the Bocas del Toro Archipelago monitored from 1999-2004 document an annual mean seawater temperature of 28.5°C, ranging from a mean of 25.9°C in Jan-Feb to 29.7°C in Sept-Oct (Kaufmann and Thompson, 2005).

**Figure 2.**
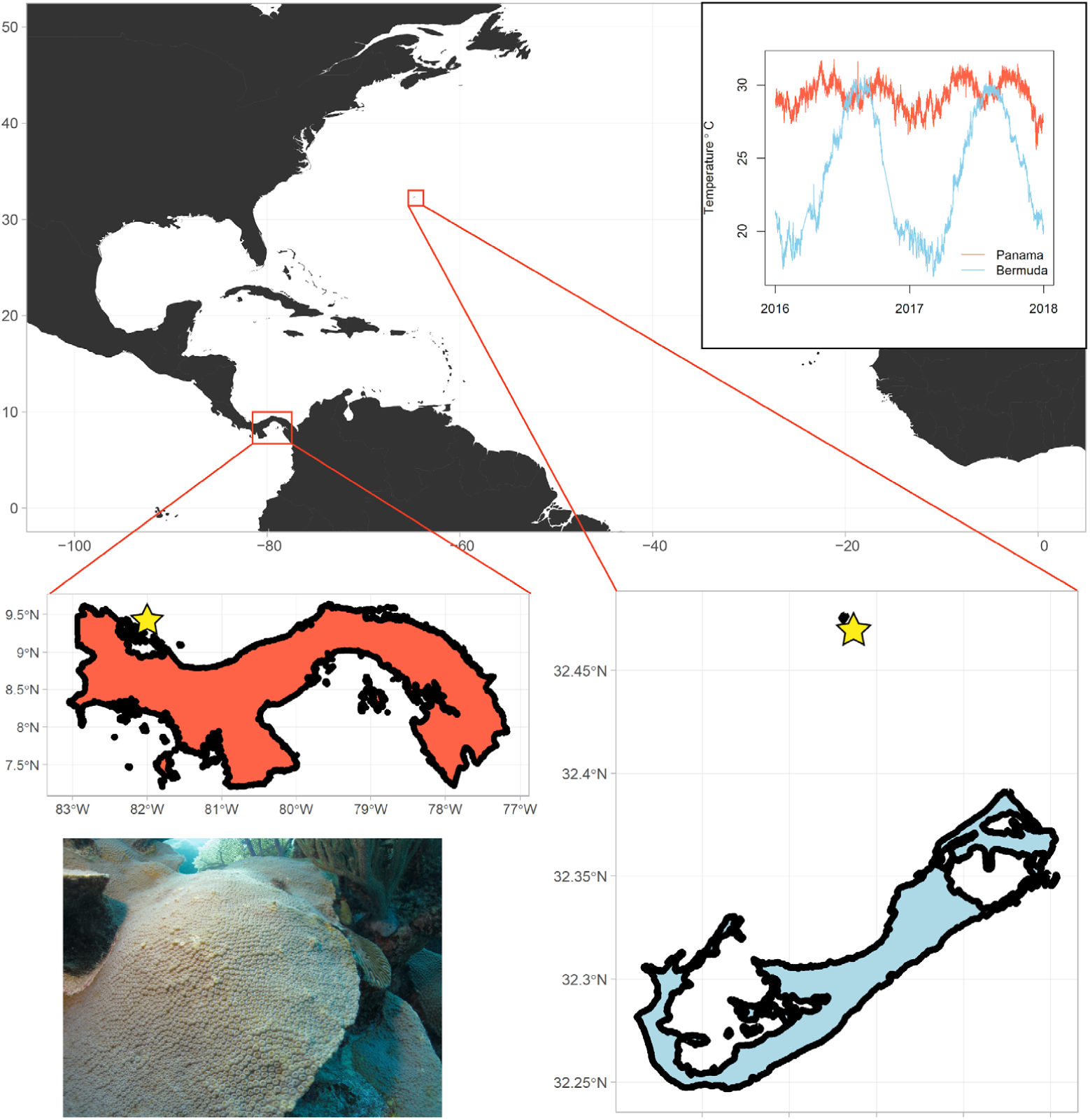
Map of study sites, thermal histories, and image of *Orbicella franski*. Yellow stars in the Panama (orange) and Bermuda (blue) maps indicate collection sites. Inset in the top right shows the thermal histories for each location (Data sets provided by the Physical Monitoring Program of the Smithsonian Tropical Research Institute; NOAA National Data Buoy Center) and inset on the bottom left is an image of *O. franksi* from the collection site in Bermuda (PC: N. Silbiger).

Located at approximately 32°N, 64°W, 1049 km (652 nm) southeast of Cape Hatteras (US central east coast), Bermuda’s sub-tropical coral reefs represent the northernmost shallow-water reef system in the Atlantic Ocean (Figure 2). Annual reefal temperatures across the shallow reef platform (3–18 m depth) range from 15–30°C (Coates et al., 2013; Locke et al., 2013a), which allows a variety of tropical marine organisms to live in this region, including 38 hermatypic and ahermatypic scleractinian coral species (Locke et al., 2013b). Bermuda is markedly cooler, however, than typical Caribbean reefs. For example, during the wintertime, on average, inshore SST is 8°C cooler in Bermuda than Panama (Figure 2). Maximum temperature in the summer is also >1°C lower in Bermuda than in Panama. Importantly, this has led to minimal impacts of coral bleaching in Bermuda (Cook et al., 1990).

### Study Species

*Orbicella* spp. has been a dominant reef building coral in the Caribbean since at least the late Pleistocene (for ~1.2 million years) (Aronson and Precht, 2001). The *Orbicella* spp. complex has a broad geographic distribution across the western Atlantic, ranging from Brazil in the south to Bermuda in the north (Budd et al., 2012) and is a vital component of Caribbean reefs. The Caribbean has warmed at a rate of 0.27°C per decade between 1985 and 2009 (Chollett et al., 2012), causing mass mortality of *Orbicella* via bleaching and infectious-disease outbreaks (e.g., Bruckner and Hill, 2009; Weil, 2004). *O. franksi* is, therefore, an ideal coral for our study given its abundance in both Panama and Bermuda, its reef-building role, and its recent listing as threatened under the U.S. Endangered Species Act.

### Sample Collection

In Bermuda, specimens of *O. franksi* were collected from the reefs at Hog Breaker, which is located on the rim reef of the northern lagoon (32° 27’ 26.38”N, 64° 50” 5.1”W; Figure 2), at a depth of 8-12m on Sept. 30, 2017. Bottom seawater temperature at the time of collection was recorded on a Shearwater Petrel dive computer as 26.1°C. In Panama, specimens of *O. franksi* were collected from the reefs at Crawl Cay on Nov. 25, 2017, which sits on the ocean facing side of the archipelago between Isla Bastimentos and Isla Popa (9° 14’ 37.8”N 82° 08’ 25”W; Figure 2), at a depth of 5-10m. This site was selected based on its distance from the mainland and the town of Bocas del Toro, in order to minimize the impacts of terrestrial runoff and nearshore anthropogenic impacts, and to more closely reflect the conditions of the rim reef collection site in Bermuda. Bottom seawater temperature at the time of collection was recorded on a Shearwater Petrel dive computer as 27.8°C. All samples were collected with a hammer and chisel. Sampled colonies in both locations were separated by a minimum of 5m to reduce the probability of selecting clones.

Samples were brought back to the respective marine laboratory (Bermuda Institute of Ocean Sciences [BIOS] or Smithsonian Tropical Research Institute [STRI]) submerged in seawater in insulated coolers. Once at the lab, the colonies were immediately fragmented with a hammer and chisel into 11 replicate ramets and maintained in seawater flow through systems outside under ambient light and ambient temperature conditions (~27°C and ~28°C in Bermuda and Panama, respectively), where they were allowed to recover for 24-72 hrs. before experimentation. Samples were then moved to a holding tank inside the laboratory in ambient seawater temperature under greenhouse lights in Bermuda (Sun Blaze T5 High Output Fluorescent Light Fixtures) at 130±6 (mean ± SE, n=9) µmol m^-2^ s^-1^ prior to TPC measurements.

### PI Curves

Prior to experimental exposures, replicate coral fragments were used to generate photosynthesis-irradiance curves for each location to determine saturating irradiance for assessing rates of photosynthesis. Fragments were placed in individual acrylic respiration chambers (620mL) with 5µm filtered seawater in Bermuda and 50µm filtered in Panama, with individual temperature (Pt1000) and fiber-optic oxygen probes (Presens dipping probes [DP-PSt7-10-L2.5-ST10-YOP]). Oxygen was measured every second in the coral chambers and blank chambers. Fragments were exposed to nine light levels generated by LED lights hung above the chambers (Arctic-T247 Aquarium LED, OceanRevive): 0, 31, 63, 104, 164, 288, 453, 610, and 747 µmol m^-2^ s^-1^ in Bermuda and 0, 22, 65, 99, 210, 313, 476, 613, and 754 µmol m^-2^ s^-1^ in Panama. Light levels were determined by an underwater cosine corrected sensor (MQ-510 quantum meter Apogee Instruments, spectral range of 389-692 nm ± 5 nm).

Rates of Oxygen flux were extracted using repeated local linear regressions with the package *LoLinR* (Olito et al., 2017) in R (R Core Team, 2013), corrected for chamber volume, blank rates, and normalized to coral surface area calculated by tracing of planar area of the flat O. franksi samples using ImageJ (Schneider et al., 2012). *LoLinR* was run with the parameters of L_pc_ for linearity metric (L_pc_ = the sum of the percentile-ranks of the Zmin scores for each component metric) and alpha = 0.2 (minimum window size for fitting the local regressions, which is the proportion of the total observations in the data set) for observations, and thinning of the data from every second to every 20 seconds. A non-linear least squares fit (NLLS;(Marshall and Biscoe, 1980)) for a non-rectangular hyperbola was used to identify PI curve characteristics of each species. This model is as follows:

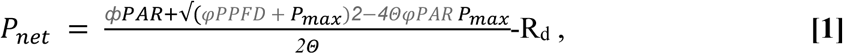

where the parameters are P_net_ and P_max_ (area-based net and maximum gross photosynthetic rates, respectively), R_d_ (dark respiration, started at min rate in the dark), AQY (φ, apparent quantum yield), PAR (photosynthetically active radiation), and Theta (Θ, curvature parameter, dimensionless).

These PI curves identified saturating irradiance (I_k_) of 110 µmol m^-2^ s^-1^ for Bermuda and 184 µmol m^-2^ s^-1^ for Panama, with no indication of photoinhibition (Figure S1). Subsequent measurements of photosynthetic rates were completed at 553±22 µmol m^-2^ s^-1^ in Bermuda and 623±21 µmol m^-2^ s^-1^ in Panama.

### Characterizing Metabolic Thermal Response

For TPC measurements, fragments were placed in individual respiration chambers to measure photosynthesis and dark respiration rate, as light enhanced dark respiration rates (Edmunds and Davies, 1988) (hereafter, respiration or R_d_), after ~60 minutes of light exposure. The respirometry setup consisted of ten 620 ml chambers with magnetic stir bars. Samples were measured in a series of runs that consisted of replicate fragments and blank chambers, and included 60 minutes under saturating irradiance, followed by 60 minutes of dark. New fragments from the same colonies (N = 4 colonies/genotypes) were used for each temperature run, resulting in acute TPC curves. Importantly, while these acute and non-ramping TPCs provide good comparative (relative) metrics of thermal responsiveness, they overestimate the metrics relative to samples acclimatized to each temperature, or ramped through all the temperatures (Schulte et al., 2011; Sinclair et al., 2016). We measured net photosynthesis and dark respiration at seven temperatures in Bermuda (24, 26, 27, 29, 31, 32, 34, 36°C) and eleven temperatures in Panama (26, 27, 28, 29, 30, 31, 32, 33, 34, 35, 37°C). Temperature was controlled to ±0.1°C by a thermostat system (Apex Aquacontroller, Neptune Systems) using a chiller (AquaEuroUSA Max Chill-1/13 HP Chiller) and heaters (AccuTherm Heater 300W). Rates of oxygen flux were extracted following the methods described above and gross photosynthesis (GP) was calculated as the absolute values of net photosynthesis plus dark respiration.

In Bermuda only, we also measured light and dark calcification across the seven temperatures. Calcification rates were calculated using the total alkalinity (A_T_) anomaly technique (Chisholm and Gattuso, 1991). Water samples (N = 3 replicates) for A_T_ were collected in thrice rinsed, acid washed 250 mL Nalgene bottles from the temperature-controlled seawater prior to incubation and then again from each chamber (both corals and blanks) after the 60 min incubation. A_T_ samples were immediately preserved with 100 µL of 50% saturated HgCl_2_. A_T_ was analyzed using open cell potentiometric titrations (Dickson et al., 2007) on a Mettler T5 autotitrator. A certified reference material (CRM, Reference Material for Oceanic CO_2_ Measurements, A. Dickson, Scripps Institution of Oceanography) was run at the beginning of each sample set. The accuracy of the titrator was always less than 0.8% off from the standard and the precision was <5 µmol kg^-1^ between sample replicates.

Because calcification from the alkalinity anomaly is the sum of all calcification and dissolution processes in the coral, all exposed skeleton on the corals was covered with parafilm immediately prior to measurements to minimize dissolution of the carbonate framework. Light and dark calcification rates (µmol CaCO_3_ cm^-2^ h^-1^) were calculated using the following equation:

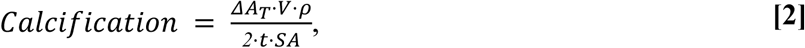

where ΔA_T_ (µmol kg^-1^) is the difference in A_T_ between the initial and post incubation sample (note: ΔA_T_ in the blanks was subtracted from the ΔA_T_ in the coral samples to account for any calcification due to other calcifiers in the seawater), V (cm^3^) is the volume of water in the chamber accounting for the volume of the coral, ρ is the density of seawater (average density = 1.023 g cm^-3^), t (h) is the incubation time (~1 hr), and SA (cm^2^) is the surface area of the corals determined tracing of planar area of the flat *O. franksi* samples using ImageJ (Schneider et al., 2012). ΔA_T_ was divided by 2 because 1 mol of CaCO_3_ is produced for every 2 mols of A_T_. Changes in dissolved inorganic nutrients were assumed to be minor in an hour incubation, making it unnecessary to account for nutrient concentrations in the alkalinity anomaly. Salinity was also measured in the pre-and post-incubation water samples (~37 psu), but no evaporation was noted: the chambers were airtight. We present our results as net calcification (NC = light calcification + dark calcification).

### Model Construction

We used Bayesian hierarchical models with Markov Chain Monte Carlo (MCMC) simulations to estimate coral thermal tolerance metrics for GP, R_d_, and NC. One outlier in the NC data (temperature = 26 °C) was removed due to contamination of an alkalinity sample. Log(x+1) GP, R_d_, and NC rates were fit to modified Sharpe-Schoolfield models for high temperature inactivation (Schoolfield et al., 1981) (Table 1, Fig 1). We ran three separate models to explicitly test the hypotheses that TPC parameters for GP (model 1) and R_d_ (model 2) differ between the Bermuda and Panama populations, and that TPC parameters differ among the three organismal functions in Bermuda only (model 3). To get reliable estimates of all TPC parameters, the maximum experimental temperature needs to be high enough to bring the measured rate to near zero. We did not achieve near-zero R_d_ rates in Panama (highest temperature measured was 37 °C), thus making the Panama versus Bermuda comparison for R_d_ unreliable. Therefore, for comparisons between populations (Panama versus Bermuda), we only present the GP model in the main text and we included the R_d_ model results in the supplement (see Figures S2-4). T_opt_ and CT_max_ were both estimated within the MCMC chain. CT_max_ was calculated as the temperature at which there was a 90% loss of maximum rate (i.e. rate at T_opt_). For detailed model description please see supplemental materials.

### Model Fitting and Analysis

We ran our model using MCMC algorithms in JAGS (just another Gibbs sampler) (Plummer, 2003) called from R (R Core Team, 2013) using the R packages, *rjags* (Plummer, 2011) *and dclone* (Sólymos, 2010). We ran three parallel chains of length 2.5M, with a burn-in of 2M, and a thinning parameter of 1/2000 to account for high autocorrelation in the chains, leaving a total of 13,500 samples for inference.

We assessed convergence by checking all trace plots, ensuring that all chains were well-mixed, and calculating Gelman-Rubin statistics (Gelman and Rubin, 1992) for all parameters (all of which were < 1.05). To assess model fit, we used posterior predictive checks by adding a step in each MCMC iteration to simulate data based on our model’s posterior predictive distribution and then comparing it to our observed dataset. Goodness of fit was evaluated using Bayesian p-values, which are based on comparing the discrepancies between observed and simulated data. Bayesian p-values for the mean, standard deviation, and coefficient of variance for all models were between 0.49 - 0.53 (close to 0.5), indicating that differences between observed and simulated data are likely due to chance. Lastly, we plotted our observed versus predicted data from the model simulations and they were in close agreement (Fig S5-6).

For our numerically generated posterior samples, we report median values with two-tailed 95% Bayesian credible intervals (BCI) for each parameter (essentially, Bayesian confidence intervals). We used the *compare_levels* function in the *TidyBayes* package (Kay, 2018) to make pairwise comparisons of each parameter among populations and organismal functions. Pairwise comparisons with credible intervals that do not overlap zero are considered to be statistically different from each other. All R and JAGS code and data will be available on github (https://github.com/njsilbiger/CoralThermalTolerance) and citable at Zenodo upon publication.

## Results

### Differences in TPC parameters between populations

The Panama and Bermuda coral populations had markedly different functional responses to temperature (Figure 3). Specifically, for GP, the corals from Panama were more thermally tolerant than those from Bermuda, with a 2.17°C higher T_opt_ (0.95 - 3.65°C [95% BCI]) and a 1.59°C higher CT_max_ (0.93 - 2.61 °C [95% BCI]; Figure 4; Figure S7). The Panama population also had higher GP rates overall. At the reference temperature (27°C) the log(x+1) GP rate was 0.24 µmol cm^-2^ hr^-1^ higher (0.13 - 0.32 µmol cm^-2^ hr^-1^ [95% BCI]) in Panama than Bermuda (Figure 4; Figure S7). Panama corals also had a marginally steeper deactivation energy (E_h_) than Bermuda corals, meaning GP drops out more quickly in Panama once it reaches its thermal optimum, but the activation energy (E) was the same between the two populations (Figure 4). For dark respiration comparisons between Panama and Bermuda see supplemental material (Figures S2-4).

**Figure 3.**
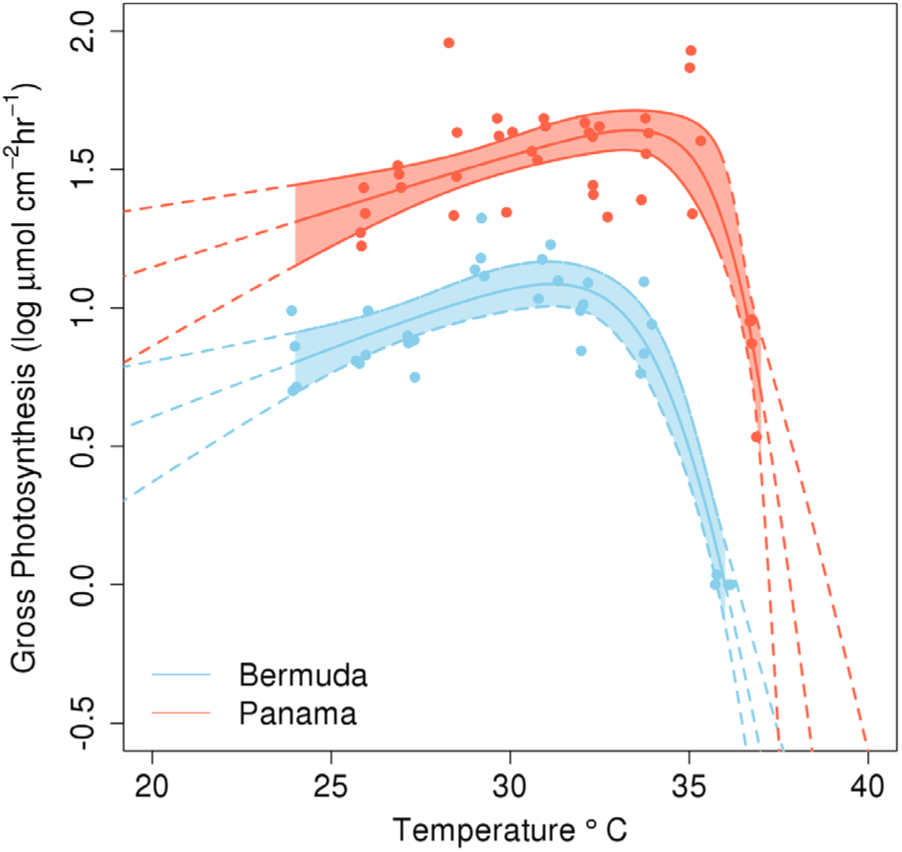
Thermal performance curves of log(x+1) gross photosynthesis rates (µmol O_2_ cm^-2^ hr^-1^) from Panama (orange) and Bermuda (blue). Each dot represents an individual fragment (n = 28 in Bermuda; n = 44 in Panama) of *Orbicella franski* from 4 putative clones. Lines are medians ± 95% BCI drawn from the posterior distribution. The shaded regions are the temperatures where data were collected.

**Figure 4.**
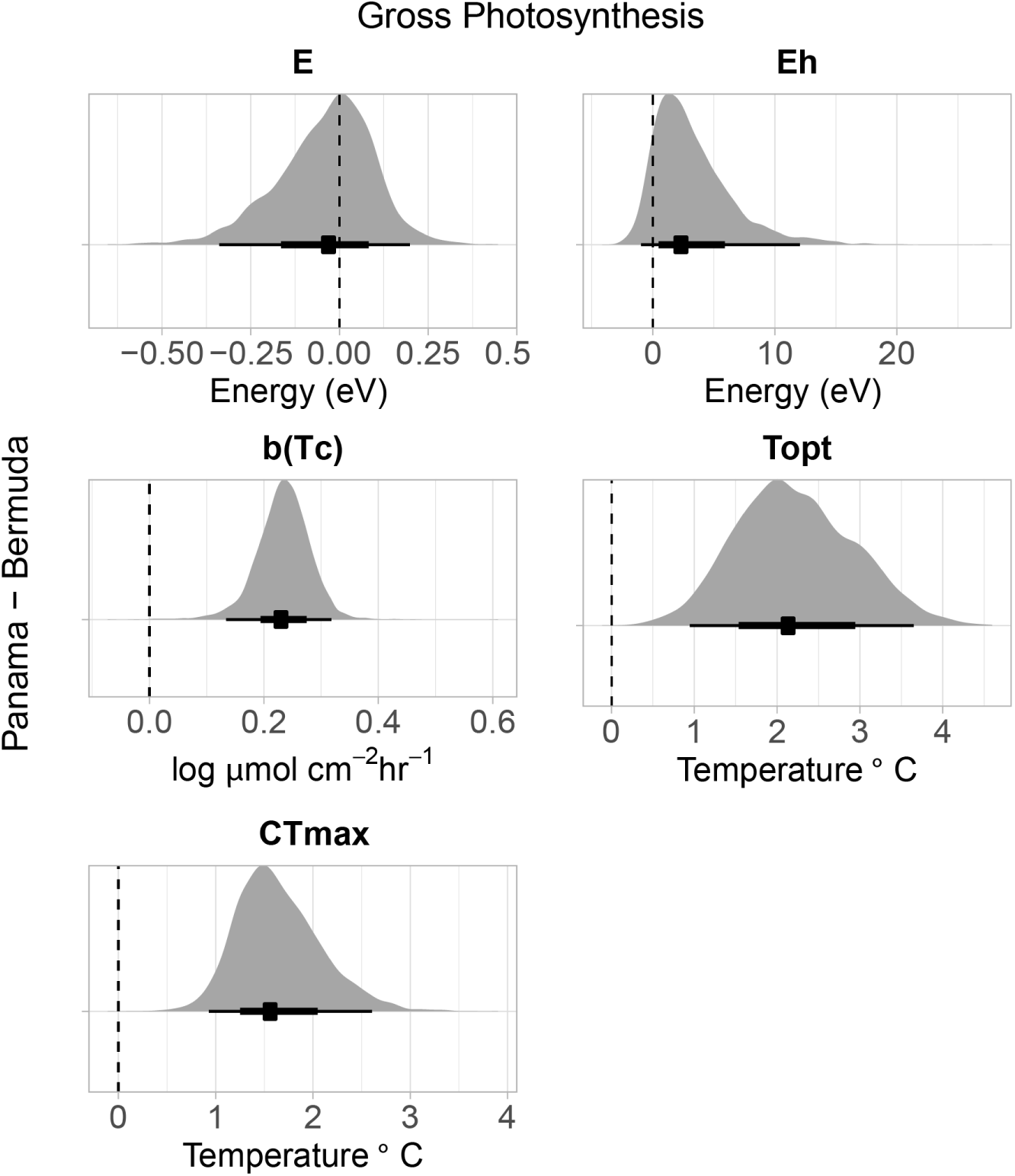
Pairwise comparisons of thermal performance metrics from Panama and Bermuda for gross photosynthesis. Dot and whiskers below each distribution are the median and 95% BCI for each parameter. If the whiskers do not cross the dashed vertical line at 0 then the populations are considered to be statistically different from one another. Positive differences indicate the metric is higher in Panama than Bermuda corals.

### Differences in TPC parameters among organismal functions within a population

The TPCs among the three organismal functions tested (GP, R_d_, and NC) were also substantially different from one another (Figure 5). In Bermuda, the general pattern in thermal tolerance was R_d_ > GP > NC (Figure 6; Figure S8). Specifically, R_d_ had the highest T_opt_ (Figure S8) and was 0.67°C (−0.4 - 1.72 °C [95% BCI]) higher than GP and 1.69°C (0.72 - 3.02°C [95% BCI]) higher than NC (Figure 6). While none of the CT_max_ values were statistically different from one another, on average R_d_ still had the highest CT_max_ (Figure S8). GP had the steepest deactivation energy (E_h_) and was 3.15 (1.36 −5.27 [95% BCI]) and 2.92 (0.89 - 5.07 [95% BCI]) higher than NC and R_d_, respectively (Figure 6). None of the activation energies (E) were significantly different from one another.

**Figure 5.**
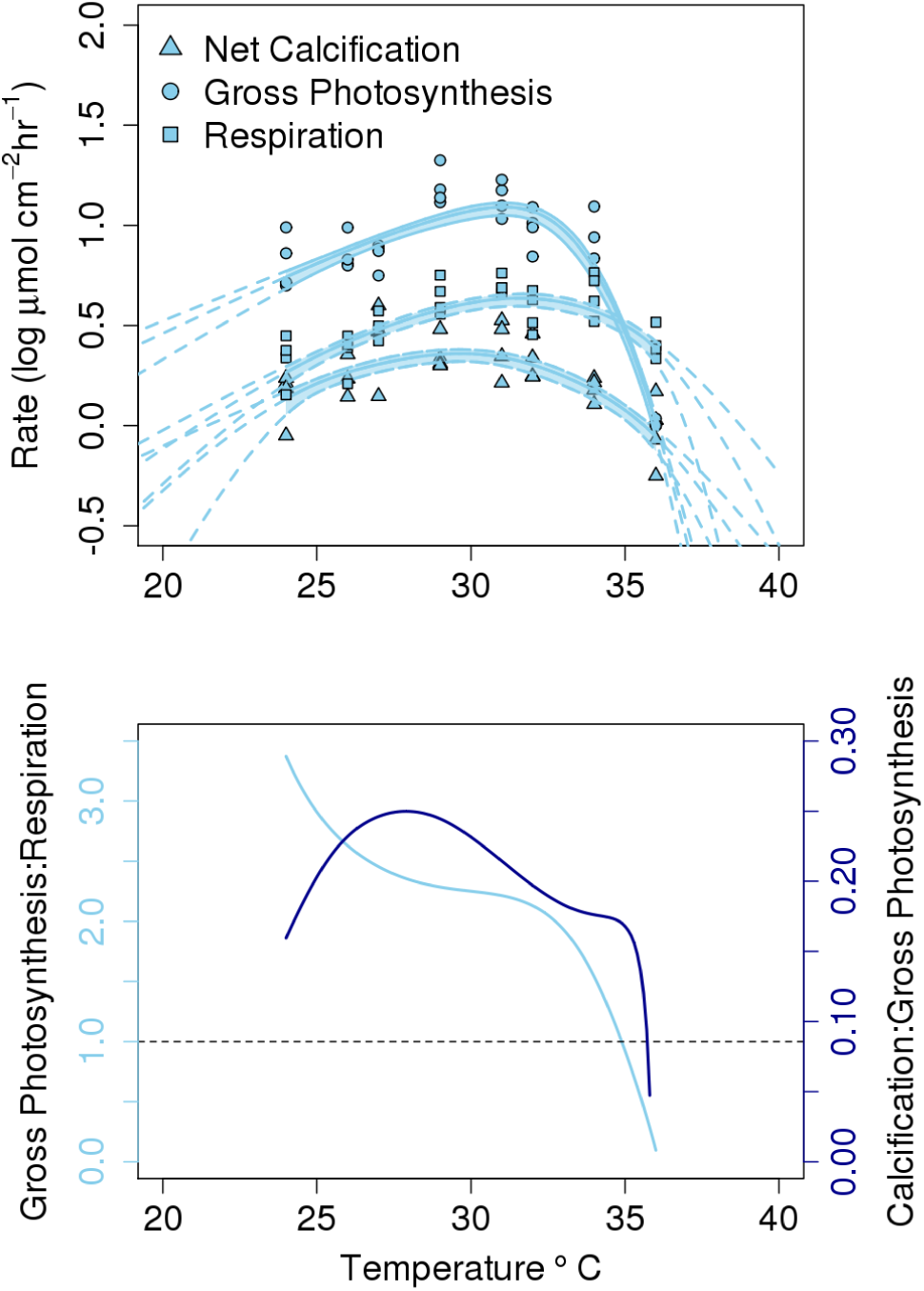
**(A)** Thermal performance curves of log(x+1) gross photosynthesis (circles), dark respiration (squares), and net calcification (triangles) rates (µmol O_2_ or CaCO_3_ cm^-2^ hr^-1^). Each dot represents an individual fragment (n = 28) of *Orbicella franski* from 4 putative clones across 7 temperatures. Lines are medians ± 95% BCI drawn from the posterior distribution. The shaded regions are the temperatures where data were collected. **(B)** GP:R (light blue) and NC:GP (dark blue) lines by temperature. Lines are from the best fit (median) from the model (a) and the rates were back transformed before taking the ratio. Dashed horizontal line is at a GP:R = 1. Any value <1 indicates that the dark respiration rate is higher than the gross photosynthesis rate.

**Figure 6.**
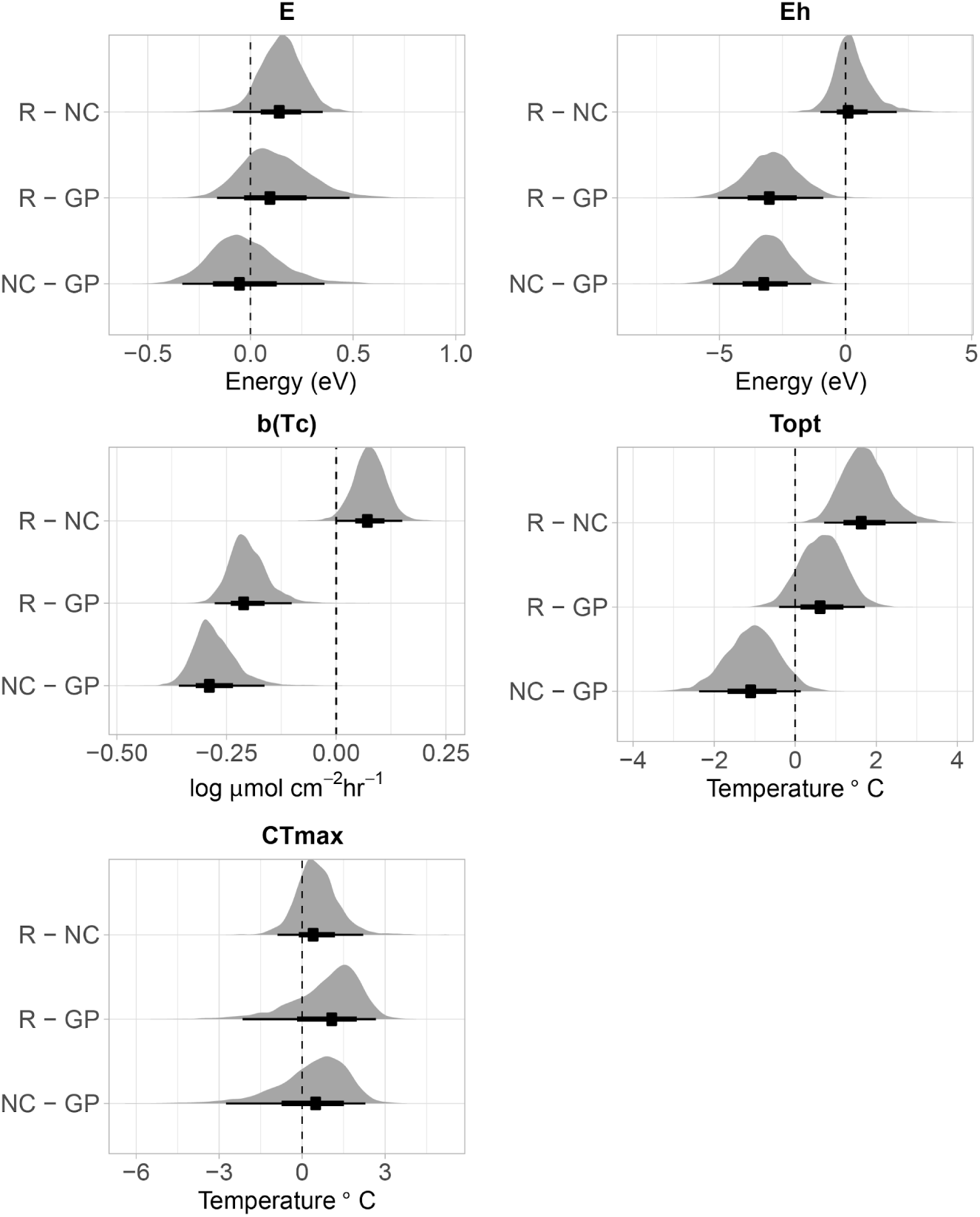
Pairwise comparisons of thermal performance metrics from Bermuda comparing the three organismal functions measured. R is light enhanced dark respiration, GP is gross photosynthesis, and NC is net calcification. Dot and whiskers below each distribution are the median and 95% BCI for each parameter. If the whiskers do not cross the dashed vertical line at 0 then the functions are considered to be statistically different from one another.

The GP:R_d_ and NC:GP ratios also varied by temperature (Figure 5b). GP:R_d_ generally declined with temperature, although it leveled off between approximately 27 and 30°C. The GP:R_d_ ratio reached 1, where gross photosynthesis and respiration were equal, at 34.9°C. NC:GP followed a unimodal curve, where the highest ratio (i.e, the most efficient calcification per unit production rate) was at 27.9°C.

### Variance components in thermal performance metrics

Clone-level variation in TPC metrics was generally low (Figure S9). Variance in b(T_c_) due to differences between populations was 3.6× higher than variance due to differences among clones within a site. Similarly, variance in b(T_c_) among the three organismal functions was 2.7× higher than variance among clones.

## Discussion

Our results indicate that populations with different thermal histories respond differently to acute warming. Specifically, corals in Bermuda were less heat tolerant than those in Panama. Both T_opt_ and CT_max_ were greater for photosynthesis and T_opt_ was also greater for respiration (note the statistical significance of this difference was marginal). These observed differences in thermal tolerance roughly match the difference in the average maximum summertime high at the two locations (Panama maximum temperature is ~1.2°C higher than Bermuda; Figure 2). Importantly, there are several differences in the thermal regimes between Panama and Bermuda, and both the mean and variance in environmental temperature can affect coral metabolism (Putnam and Edmunds, 2011) and thermal sensitivity (e.g., Safaie et al., 2018). While there are several other environmental differences between Bermuda and Panama, a likely explanation for the observed thermal sensitivities is the fairly large (for a tropical and sub-tropical system) differences in temperature (Figure 2). However, the comparison of phenotypic traits between populations in this, or any other latitudinal study, should be interpreted with caution as the results cannot be clearly attributed to temperature alone.

Assuming the local temperature regime is a dominant cause of the observed among-population differences, numerous evolutionary and ecological processes could underlie this environmental matching. First, natural selection for thermally tolerant host genotypes in Panama could lead to adaptation to the local thermal regime and genetic population differentiation (Torda et al., 2017). Second, both the coral host and endosymbiont could be physiologically acclimatized to local temperatures (Brown and Cossins, 2011). Third, numerous epigenetic mechanisms could enable the Panama corals to be more tolerant of extreme high temperatures (Eirin-Lopez and Putnam, 2019). Fourth, the dominant symbionts could be genetically differentiated (Baker et al., 2004), with a more thermally tolerant strain present at the Panama site. Numerous studies have found substitutions of thermally sensitive endosymbionts by more tolerant ones in both space and time (e.g., Baker et al., 2004; Boulotte et al., 2016). Fifth, it is also possible that differences in the coral-associated microbial community affect responses of host and symbiont to the temperature treatments (Webster and Reusch, 2017). While our study was not designed to tease apart the relative contributions of these or other potential mechanisms leading to less thermally sensitive genotypes at the warmer site, several studies have found evidence supportive of thermal acclimation across spatial gradients (Howells et al., 2012; Sunday et al., 2011).

In addition to differing thermal sensitivity among populations, we also found that there are differences in thermal sensitivity among metabolic processes within populations. Specifically, GP, R_d_, and NC all had different thermal optima (T_opt_), rates of deactivation (E_h_), and rates at a reference temperature (b(T_c_)) (Figure 6). Different enzymatic machinery is utilized for each of these traits (e.g., citrate synthase in respiration, RuBISCo in photosynthesis, and Ca-ATPase in calcification), with some being specific to host or symbiont function. Therefore, in comparison, we hypothesized *a priori* that holobiont respiration was likely to be less sensitive to temperature than photosynthesis, which was supported by our data. Calcification was the least tolerant trait with the lowest thermal optimum. Other coral studies have also demonstrated that calcification is more sensitive to temperature than photosynthesis and respiration (though not using an explicit TPC approach) (e.g., Al-Horani, 2005; Reynaud et al., 2003). For example, at high temperatures, *Galaxea fascicularis* in the Red Sea produced less O_2_ than it consumed (i.e. higher respiration than photosynthesis rate) and began to decalcify (Al-Horani, 2005). Together, these results indicate that corals may be able to survive slight increases in warming (e.g., ~1-2°C), but they would still experience declines or even ecological loss of important functions related to fitness, or that are necessary for coral reef ecosystem functioning, such as net ecosystem production and net ecosystem calcification.

The ratios of GP:R_d_ and NC:GP both varied with temperature (Figure 5). These ratios and how they change with temperature, have implications for survival and the long-term persistence of certain functions. For example, corals need a GP:R_d_ ratio of greater than 2 to maintain long-term autotrophy (Coles and Jokiel, 1977). Here, we saw GP:R_d_ generally declined with temperature, a pattern that has been shown in other coral studies (Coles and Jokiel, 1977), and that the GP:R_d_ dropped below 2 at 32.7°C. Therefore, as ocean temperatures continue to warm, *O. franski* will need to shift to more heterotrophic food sources to survive. The NC:GP ratio had a unimodal response to temperature, with the optimal calcification per unit production rate at 27.9°C. Calcification and photosynthesis are highly coupled in corals and calcifying macroalgae (Barnes and Chalker, 1990; Gattuso et al., 1999; Goreau, 1959; Schneider and Erez, 2006), although the mechanisms linking them continue to be debated (see, Gattuso et al., 1999; Gattuso et al., 2000). Assessing NC:GP ratios can uncover the amount of CO_2_ that can potentially be supplied from calcification to photosynthesis (Gattuso et al., 1999) and how this relationship may change with temperature.

We measured acute TPCs, which can be thought of as an instantaneous thermal “stress test”. While short acclimation times such as ours would underestimate absolute acclimation potential of the tested organisms, as they do not have time to fully acclimate, acute TPCs are used widely for comparative analyses (Schulte et al., 2011). For example, a relatively recent database compilation of studies examining thermal performance contains thousands of entries for >200 traits across taxa ranging from microbes to animals, spanning ~16 orders of magnitude in body size extracted from ~300 studies (Dell et al., 2013). These TPC studies have helped uncover constraints on thermal acclimation (Rohr et al., 2018) and develop critical advancements in theory (e.g., Metabolic Theory of Ecology; Brown et al., 2004). Such TPC approaches, when conscious of important assumptions and applying appropriate experimental frameworks (Schulte et al., 2011; Sinclair et al., 2016), can provide useful metrics of comparison between organisms, populations, and species.

While there is a massive body of literature on thermal tolerance of terrestrial and marine organisms (Dell et al., 2011), there is surprisingly less on the thermal performance characteristics of coral reef organisms in an explicit TPC context (but see, Aichelman et al., 2019; Jokiel and Coles, 1977; Jokiel and Coles, 1990; Rodolfo-Metalpa et al., 2014). Understanding thermal performance of corals is essential for projecting coral reef futures, given that key biological functions necessary to sustain coral reef ecosystems (e.g., photosynthesis, respiration, calcification) are thermally-mediated. Here, we advance this area of research by applying the TPC approach to a variety of fitness related parameters in a reef building coral across its geographic range, in a robust statistical framework. We suggest that future studies should incorporate multiple species and representatives from other functional groups in order to better predict ecosystem-level responses to temperature. Understanding how patterns of response differ among genotypes will improve our understanding of the inherent variability in thermal tolerance that exists within and among species and therefore potential cascading effects of biodiversity and genetic diversity loss. Taken together, these approaches will provide information critical to informing evidence-based management and conservation of these threatened ecosystems in the face of a warming ocean.

## Author Contributions

- NJS, JB, GGG, and HP designed the experiment, collected data in Bermuda, provided materials, and edited the manuscript;
- JB and GGG collected data in Panama;
- NJS and HP processed the data;
- NJS statistically analyzed the data.

## Acknowledgements

We thank BIOS and STRI for facilities support. A. Chequer, D. Becker, E. Strand, K. Gould, J. Mata-lopez, and several students in the Northeastern University Three Seas Program helped collect and process data. J Yoder provided access to a server for running the Bayesian analyses. This is CSUN marine biology contribution# XXXX.

## Data Availability

All R and JAGS code and data will be available on github (https://github.com/njsilbiger/CoralThermalTolerance) and citable at Zenodo upon publication.

